# Borg3 controls septin recruitment for primary cilia formation

**DOI:** 10.1101/2024.06.04.597321

**Authors:** Janik N. Schampera, Friederike Lehmann, Ana Valeria Meléndez, Carsten Schwan

## Abstract

Septin GTPases form linear hexa- or octameric rods that polymerize into higher order structures. They are incorporated into the cytoskeleton and involved in vital cellular functions. Among these, they play a role in the formation of primary cilia. Primary cilia are evolutionary conserved cellular signaling hubs. While it is accepted that septins localize to primary cilia and are involved in their formation and function, the regulation of septin assembly in the confined ciliary compartment remains elusive. Here we show, that Borg3, also known as Cdc42 effector protein 5 (Cdc42EP5), is an essential component of primary cilia. Specific Borg3 localization is facilitated by switching the Rho-GTPase Cdc42 between an inactive- and active state at the base of the primary cilium. The active form of Cdc42 has a defined localization restricted to the base of the primary cilium. Knockout of Borg3 as well as dysregulation of Cdc42 reduces septin dynamics at cilia and consequently, the formation of cilia.

The study demonstrates that Borg3 is a novel and essential regulator of ciliogenesis through the spatiotemporal control of septin dynamics downstream of Cdc42.

## Introduction

Primary cilia are antenna-like protrusions found on the surface of nearly every eukaryotic cell. They are non-motile structures involved in signal transduction and sensing of the cell environment playing a key role in development of organs to whole organisms (1). Dysfunction of primary cilia and cilia-associated proteins are causally related to numerous diseases, including Bardet-Biedel syndrome (BBS), Meckel-Gruber syndrome (MKS), nephronophthisis (NPHP) and polycystic kidney disease (2, 3).

Septins (SEPT) are highly conserved, small GTP-binding proteins of 30-65 kDa. In humans 13 different septins are encoded, which are categorized into four groups based on sequence homology. Septins form hexameric or octameric building blocks. These hetero-oligomers or rods can associate to form higher order structures like filaments, rings and cage-like formations that often associate with other components of the cytoskeleton (i.e. microtubules and actin filaments). Moreover, septins have a prominent role in cell signaling and membrane organization (4).

The process of ciliogenesis has been linked to septin function. First, septins were shown to form a ring-like structure at the base of cilia (5), which might act as a diffusion barrier. However, non-ring-like septin localizations at the ciliary base or along the ciliary axoneme were observed more frequently. Therefore, other studies have attributed different functions to cilia-associated septins. Septins regulate ciliary length (6) and contribute to ciliary stability by balancing the length of the distal tip (7). Moreover, important scaffolding functions of septins have been observed at the ciliary transition zone. Septin-dependent localization and interactions were observed for several transition zone proteins like Cc2d2a, B9D1, Tmem231 (8) or DZIP1L (9). Recently, it was shown that SEPT9 regulates the recruitment of transition zone proteins on Golgi-derived vesicles by activating the exocyst via the ARHGEF18 and RhoA (10). Most studies concur that inhibition of septins by any means results in less formation of cilia (5, 6, 10). While steadily important functions of septins at cilia emerge, the regulation of septin accumulation and assembly at cilia remains obscure.

Binder of Rho GTPases (Borg) proteins, also called Cdc42 effector proteins (Cdc42EPs) were not only shown to interact with septins, they are also capable of regulating their function (11). Our previous studies suggest that the Rho GTPase Cdc42 and its effector proteins Borg regulate septin assembly during the formation of toxin-induced protrusions (12) and during the formation of septin barriers against *P. aeruginosa* cellular uptake (13). Studies found that expression of constitutively active or -inactive Cdc42 as well as Cdc42-binding deficient Borg2 induces disassembly of filamentous septin structures that coalign with actin fibers in melanoma cells and cancer-associated fibroblasts (11, 14, 15). Thus, there is a connection between Cdc42, downstream Borg proteins, septins and changes in actin organization. These processes are located in the cytosol and the actin- and septin cytoskeleton mutually influence each other (16). This raises the question how septin accumulation and assembly into higher order structures is regulated in the confined compartment of the primary cilium. To this end, we study the contribution of Borg proteins to localization and dynamics of septins at primary cilia. We find that specifically Borg3 is involved in septin accumulation at cilia and efficient formation of cilia. We demonstrate that the Rho-GTPase Cdc42 facilitates Borg3-driven septin assembly by a GTPase-dependent switch that recruits Borg3 and septins to the ciliary compartment.

## Results

### Septins localize to cilia

Different functions and localizations were assigned to septin assemblies at cilia (5, 6) reflecting the tight linkage between protein localization and function. Therefore, we first analyzed the localization of septins at the cilium in Madin-Darby canine kidney cells (MDCK) and mouse inner medullary collecting duct (IMCD) cells, which are frequently used systems in studying the formation of primary cilia. Immunofluorescence studies in both cell types stained for a common member of each septin group (i.e. SEPT2, -3, -6, or -7 group) revealed that septins preferentially localize throughout the whole length of the cilium as well as in a spot-like accumulation more restricted to the base of the cilium (Figure 1 A). ∼70% of cilia had septin association along the axoneme, in ∼30% of cilia septins were restricted to the base (transition zone, TZ) and only ∼5% of cilia had no detectable septin association. This was consistent in both cell lines and resembles the results of previous studies (6, 7, 9). Representative super resolution images for both localizations are shown in Figure 1 B (along axoneme) and Figure S1 A and B (ciliary base, arrows). A ring-like localization at the base of cilia (5) was the rarest septin assembly in our experiments. Occasionally, we observed structures that resembled not fully closed rings (representative super resolution image in Figure S1 C, arrow). However, the formation of septin rings might be associated with specific stages of ciliary development or decay. The presence of a septin from each group suggests that hetero octameric protomers are present in cilia which are increasingly considered as the physiological building blocks of septin filaments which in turn exert the biological functions of septins (17).

**Figure 1.**
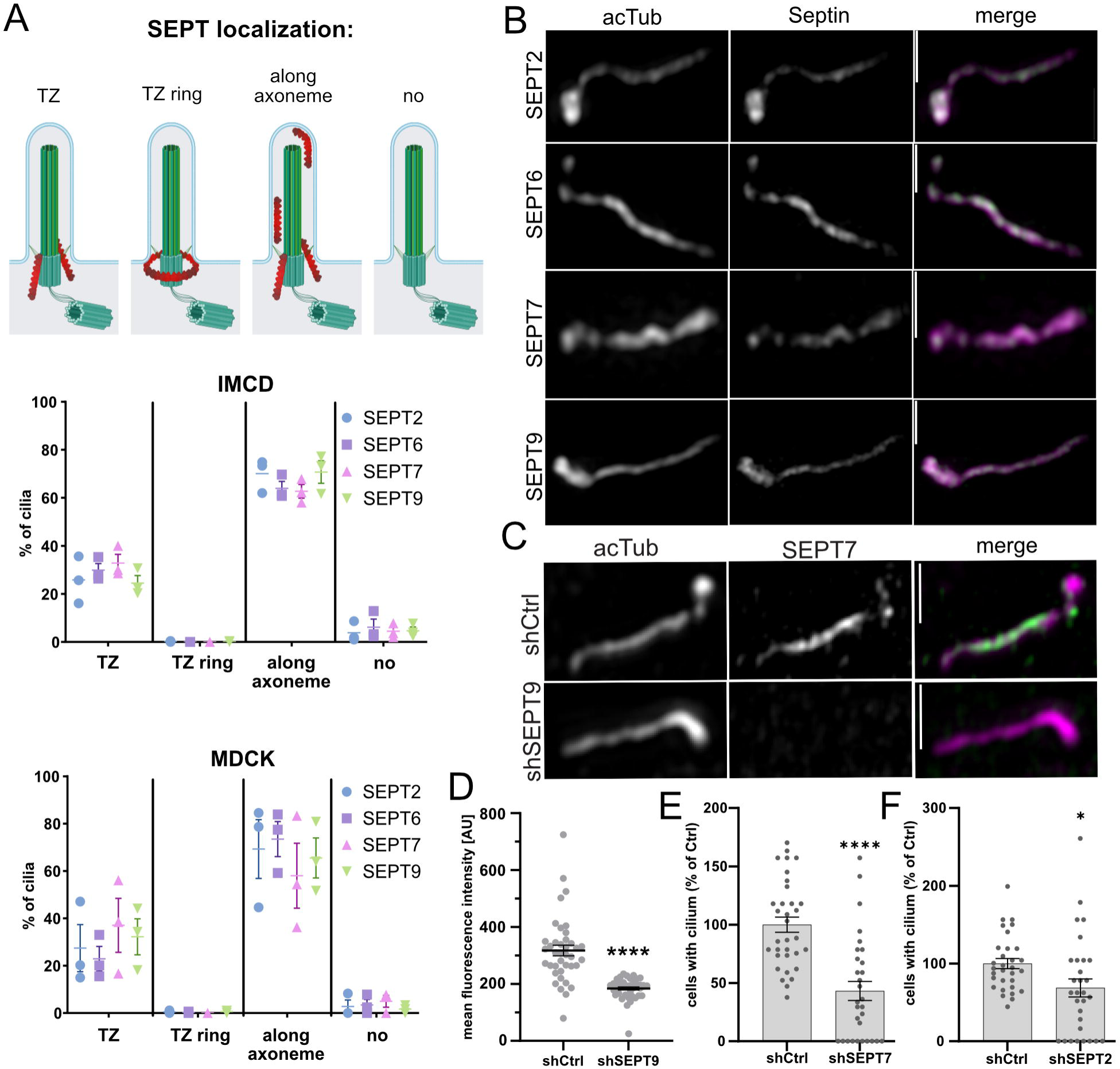
Septin localizations in primary cilia. **A)** Classification and quantification of different septin localizations at primary cilia. TZ (transition zone): Septins only found at the base of the primary cilium. TZ ring: Septins form a ring at the base of the primary cilium. Along axoneme: Septins were found throughout the length of the primary cilium. No: No Septins were found at the primary cilium. The illustration was created with BioRender. Cells were grown for 48h under starving conditions. A random panel of 9 images was acquired at 40x magnification and all observed primary cilia were classified into one of four groups regarding their septin immunofluorescence in IMCD and MDCK cells. Data are given as % of total cilia observed, mean ±SEM. n=3 **B)** Representative SIM image of IMCD cells immunostained for endogenous acetylated tubulin and SEPT2, -6, -7 or -9. Cells were serum starved for 48h before fixation. Scale bars 1 µm. **C)** Representative SIM image of MDCK cells immunostained for SEPT7 and acetylated tubulin. Cells expressed shRNA targeting SEPT9 or a non-targeting control shRNA and were starved for 48h before fixation. Scale bar 1 µm **D)** SEPT7 mean fluorescence intensity in cilia of MDCK cells treated as in C. Acetylated tubulin was used as mask for intensity measurements (AU = arbitrary units). Data are given ±SEM. >40 cilia analyzed. **E+F)** MDCK cells were transfected with non-targeting shRNAs or shRNAs targeting SEPT7 or SEPT2, respectively. Cells were starved for 48h, fixed and stained for acetylated tubulin by immunofluorescence and with DAPI to visualize nuclei. GFP additionally expressed from the shRNA vector was used to identify KD cells. The percentage of cells with a cilium was quantified. The average of the controls was set to 100% and used for normalization. Data are given as mean ±SEM. ≥30 random fields of view were analyzed from n=3.

Knockdown (KD) of SEPT9 (Figure S 1D) which diminishes the formation of functional octamers reduced septin accumulation in the ciliary compartment (Figure 1 C+D) and formation of cilia (Figure S 1E). KD of SEPT7 (Figure S1 F) which reduces formation of septin hexameric and octameric building blocks as well as KD of SEPT2 (Figure S1 G) which is needed for filament elongation inhibit ciliogenesis, too. The effect was more pronounced for SEPT7, considering that SEPT7 is the sole member of the SEPT7 group. KD of SEPT2 might be partially compensated by the expression of other members of the SEPT2-group. All in all this reveals that ciliary septin function requires septin basic building blocks (SEPT7), relies on octamers (SEPT9) and suggests that septins form filaments. Yet, septin-related extra ciliary functions for example in vesicle transport (10, 18, 19) might also influence cilia number and length. However, these results corroborate previous studies that have assigned an important role in ciliogenesis to the septin cytoskeleton (5–7, 20).

### Borg proteins colocalize with septins at cilia

The family of Borg proteins were shown to bind to septins via the conserved Borg homology 3 domain (BD3) and reorganize septins within cells (21, 22). However, Borg proteins were never observed or studied in the ciliary compartment. They are still largely uncharacterized and little is known about their regulation. Therefore, we analyzed Borg localization in cilia. Borg1, -2 and -3 were previously shown to be expressed in the kidney (23). In subconfluent MDCK cells expressing GFP-tagged versions of Borg1-3, Borgs associated with cytoplasmic septin-positive filaments (Figure S1 F), previously described in different cell-types (15, 24–26). Next, we stained for endogenous Borg proteins in confluent IMCD cells (Figure 2 A-D, Movie 1). First, we observed that Borg3 had the strongest signal in the ciliary compartment compared to Borg1 and Borg2 (Fig 2 A and B). Similar to septins, structured illumination microscopy (SIM) revealed that Borg proteins mainly localize along the cilium length and to the base of cilia in a spot-like accumulation. Septins had a more even distribution along the cilium, while Borgs appear to organize in a more clustered fashion (Fig 2 C and D). Frequently, Borg3 clusters were distributed along the cilium in an almost equidistant scattering (Movie 2).

**Figure 2.**
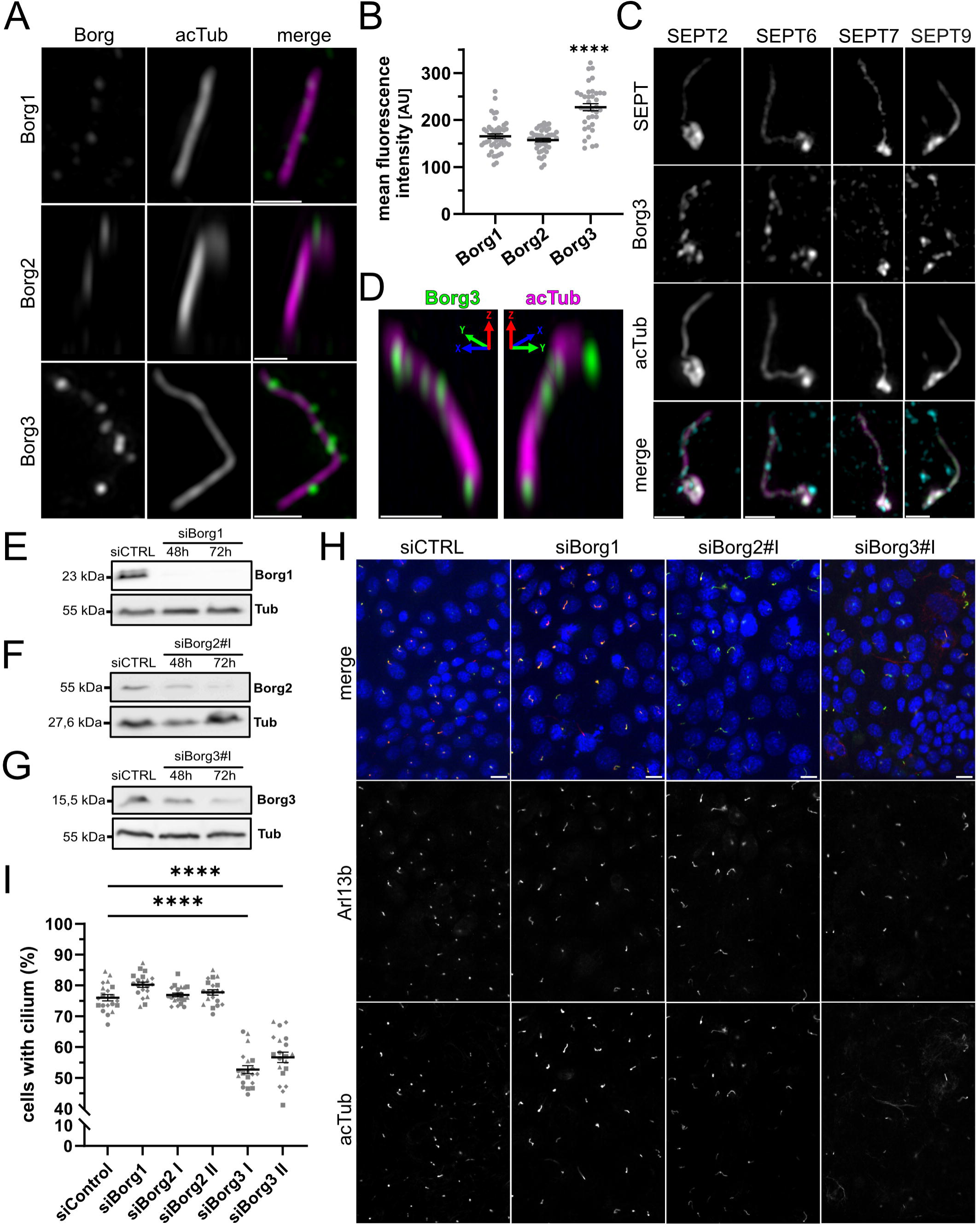
Borg3 localizes to primary cilia and is involved in cilia formation. **A)** Representative SIM image of IMCD cells immunostained for endogenous Borg1, -2 or -3 and acetylated tubulin. Cells were serum starved for 48h before fixation. Scale bar 1 µm. **B)** Quantification of Borg immunofluorescence average intensity in cilia like in A). Acetylated tubulin was used as mask for intensity measurements (AU = arbitrary units). Data are given ±SEM. ≥3 random fields of view were analyzed from n=3. **C)** Representative SIM image of IMCD cells immunostained for endogenous acetylated tubulin, Borg3 and SEPT2, -6, -7, or -9. Cells were serum starved for 48h before fixation. Borg3 and SEPT were stained with directly labeled primary antibodies. Scale bars 1 µm. **D)** Representative SIM image of IMCD cells immunostained for endogenous Borg3 and acetylated tubulin. IMCD cells were fixed in a µ-slide (Ibidi) enabling analysis of standing cilia. Image shows zoom of cilium from Movie 1. Colored arrows indicate all three dimensions with the viewing plane indicated by the horizontal and vertical arrow. Scale bar 1 µm. **E, F, G)** IMCD cells were transfected with control siRNA or with siRNAs targeting Borg1, Borg2 or Borg3 in E), F) or G), respectively. siRNA mediated KD was confirmed by western blot. α-tubulin was used as loading control (55 kDa). **H)** Representative confocal image of IMCD cells immunostained for endogenous Arl13b and acetylated tubulin. Cells were transfected as in E), F) and G) and serum starved for 72h before fixation. DAPI staining labeled nuclei. Scale bar 10 µm. **I)** Quantification of cilia formation in cells treated as in E), F), G) and H). The percentage of cells with a cilium was quantified. Data are given ±SEM. >4000 cells were analyzed from ≥25 random fields of view n=4. Symbol shapes indicate individual data points from biological replicates.

### Borg3 influences formation of cilia

In view of the specific localization of Borgs at cilia the effect of Borg depletion on cilia formation was analyzed. IMCD cells were transfected with siRNAs targeting Borg1-3. siRNA treatment for 48h resulted in downregulation of the specific Borg protein on the mRNA and protein level (Figure S1 G and Figure 2 E, F and G). KD of Borg3 by siRNA significantly reduced the number of cells developing a cilium (Figure 2 H and I). This was not observed for Borg1 and Borg2.

To further analyze the effects of Borg3 on cilia, we generated Borg3 knockout (KO) IMCD cells by CRISPR/Cas9 (27). KO analysis was performed by TIDE (https://tide.nki.nl; (28)). The inserted mutations for two generated cell lines are outlined in Figure 3 A (representative sequencing result Figure S2 A). KO#1 has a nucleotide insertion in one allele which results in a frameshift after 12 amino acids of the wildtype (wt) protein and a deletion of 2 nucleotides in the other allele resulting in a frameshift after 10 amino acids. In KO#2 both alleles carry the same mutation, an insertion of one nucleotide resulting in a frameshift after 13 amino acids. Thus, both functional domains of Borg3, the CRIB domain and the BD3 domain, are destroyed. The CRIB domain mediates the interaction with Cdc42, while the BD3 domain interacts with septins SEPT6/7 (22, 23, 29). Immunofluorescence staining of Borg3 KO#1 cell line revealed the absence of Borg3 staining in cilia (Figure S2 B).

**Figure 3.**
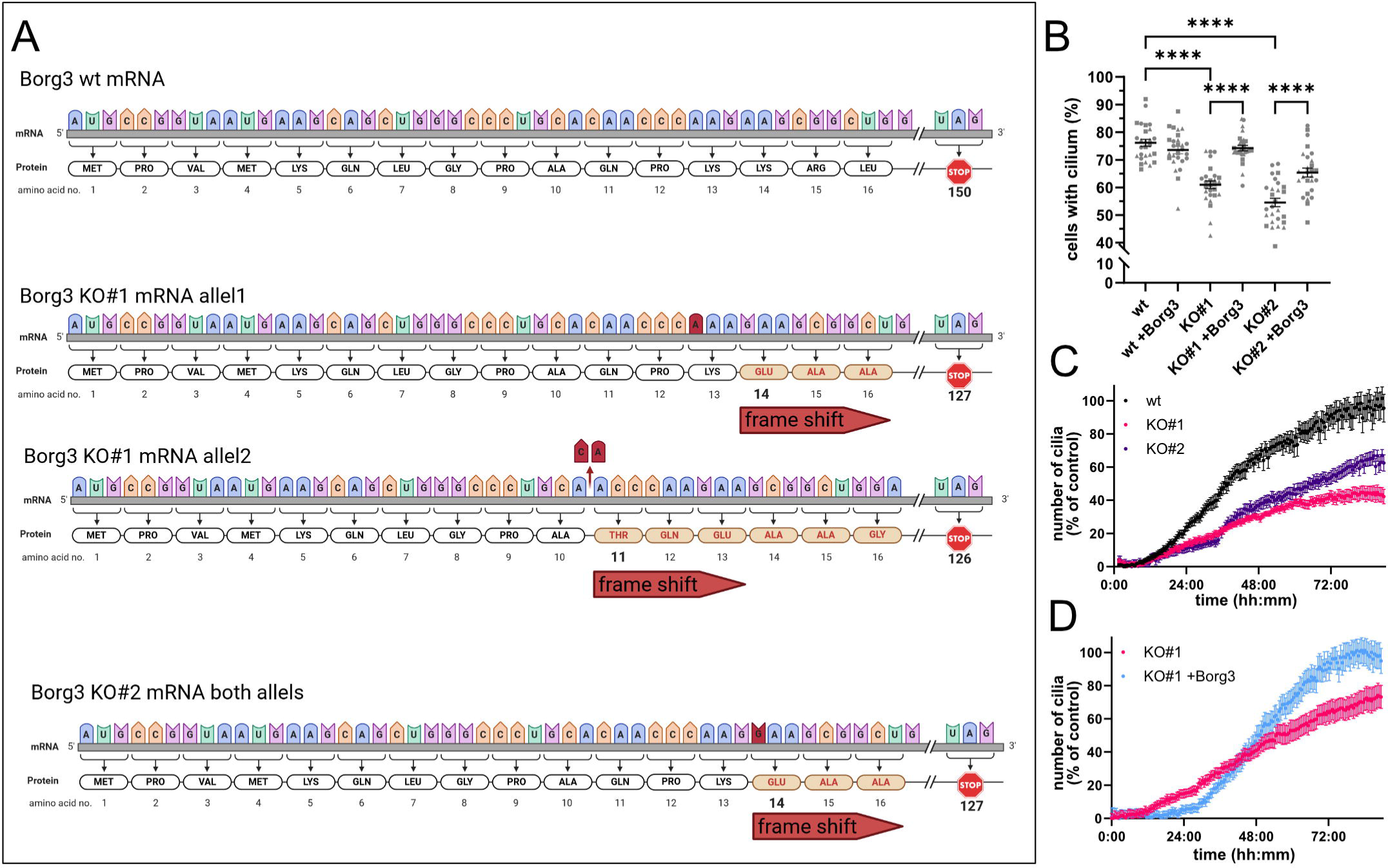
Borg3 KO inhibits formation of cilia. **A)** Schematic representation of the sequencing results from the generated CRISPR/Cas9 mediated KO of Borg3 in IMCD cells. The nucleotides lost or gained in the subsequently used KO IMCD cell lines as well as the effect on the protein translation is shown. KO cell line #1 (KO#1) has two allele specific mutations while KO cell line #2 (KO#2) has a homozygous mutation. The corresponding sequencing chromatogram is shown in Figure S2 A. **B)** Quantification of cilia formation in IMCD wt and Borg3 KO cells. Cells were grown to confluency and starved for 48h. In comparison cells transfected with GFP-Borg3 were analyzed to observe rescue effects. Cells were stained for acetylated tubulin to label cilia and with DAPI to label nuclei. Data are given as ±SEM. >4000 cells were analyzed from 9 random field of views from n=3. Symbol shapes indicate individual data points from biological replicates. **C)** Cilia growth over time in wildtype and KO IMCD cell lines stably expressing Arl13b-tomato. Cells were seeded in high and equal densities to reach immediate confluency at imaging start. Cells were observed by video microscopy for 100h with images taken every 30min. Cilia were identified by fluorescence using an intensity and size dependent filter (Figure S2 C). KO cells were normalized to the highest cilia count of the WT control. For every condition each biological replicate consisted of 3 technical replicates in which 4 random fields of view per replicate were analyzed. Data are given ±SEM. n=3 **D)** Borg3 KO IMCD cell line #1 stably expressing Arl13b was treated as in C). Additionally, KO cell line #1 stably expressing Arl13b was transfected with GFP-Borg3 1 day before the experiment and enriched by FACS as a rescue. Cells were observed and analyzed as in C). The rescue was normalized to the highest cilia count of KO cell line #1 rescue. Data are given ±SEM. n=3.

In line with the KD experiments, KO of Borg3 reduced the number of cells developing a cilium (Figure 3 B). The phenotype was rescued by transient re-expression of GFP-Borg3 in KO cells. In both KO cell lines the fraction of ciliated cells significantly increased after GFP-Borg3 expression.

Since a quantification of cilia bearing cells after fixation and immunofluorescence staining is only a snapshot of the process of cilia formation and recent studies argue that fragile parts of cilia can be easily lost during fixation and sample processing (7), we followed the formation of cilia in real time over up to 4 days. Cells stably expressing Arl13b-tomato were seeded at high density, transferred to FCS-free medium and monitored by video microscopy. These studies corroborated the immunofluorescence end-point studies, as all KO cell lines revealed a reduced and delayed formation of cilia (Figure 3 C and Figure S2 C). KO#1 cell line was rescued by transient re-expression of GFP-Borg3. After transfection cells were enriched by FACS. In cells overexpressing GFP-Borg3 the formation of cilia was increased (Figure 3 D).

### Septins at cilia are regulated by Cdc42 and its downstream effector Borg3

All Borg proteins have a conserved CRIB domain and act as downstream effectors of the small Rho GTPases Cdc42 and TC10 (11, 12, 14, 23). Therefore, it is tempting to speculate that the interplay between Cdc42 and Borgs could play a vital role in septin assembly at cilia. Moreover, previous studies suggest that Cdc42 is involved in ciliogenesis e.g., by localizing and regulating the exocyst complex (30, 31).

Confluent MDCK cells expressing 5HT6-tomato as ciliary marker were stained for SEPT2 and active Cdc42 by immunofluorescence. Active Cdc42 colocalized with SEPT2 at the base of cilia (Figure 4 A). A linescan from the ciliary base to the tip showed that the fluorescence signal of active Cdc42 was restricted to the base of the cilium. Super resolution microscopy of cells stained for active Cdc42 and Borg3 revealed that Borg3 localizes in close proximity to active Cdc42 at the base of the cilium (Figure 4 B) and also within the cilium. The Cdc42 guanine nucleotide exchange factors (GEFs) Tuba and intersectin 2 were previously identified as positive regulators of ciliogenesis and show a corresponding ciliary localization (32, 33). We observed colocalization of Tuba and Centrin2 after expression of both proteins (Figure S3 A). Expression of Cdc42 or immunofluorescence staining of Cdc42 independent of its activity state showed an accumulation at the cilium as well (Figure S3 B and C). In contrast to active Cdc42, the signal of total Cdc42 was less restricted to the base of the cilium. Cdc42 within the cilium appeared not membrane associated as we can resolve the ciliary membrane by SIM with an Arl13b staining (Figure S3 C).

**Figure 4.**
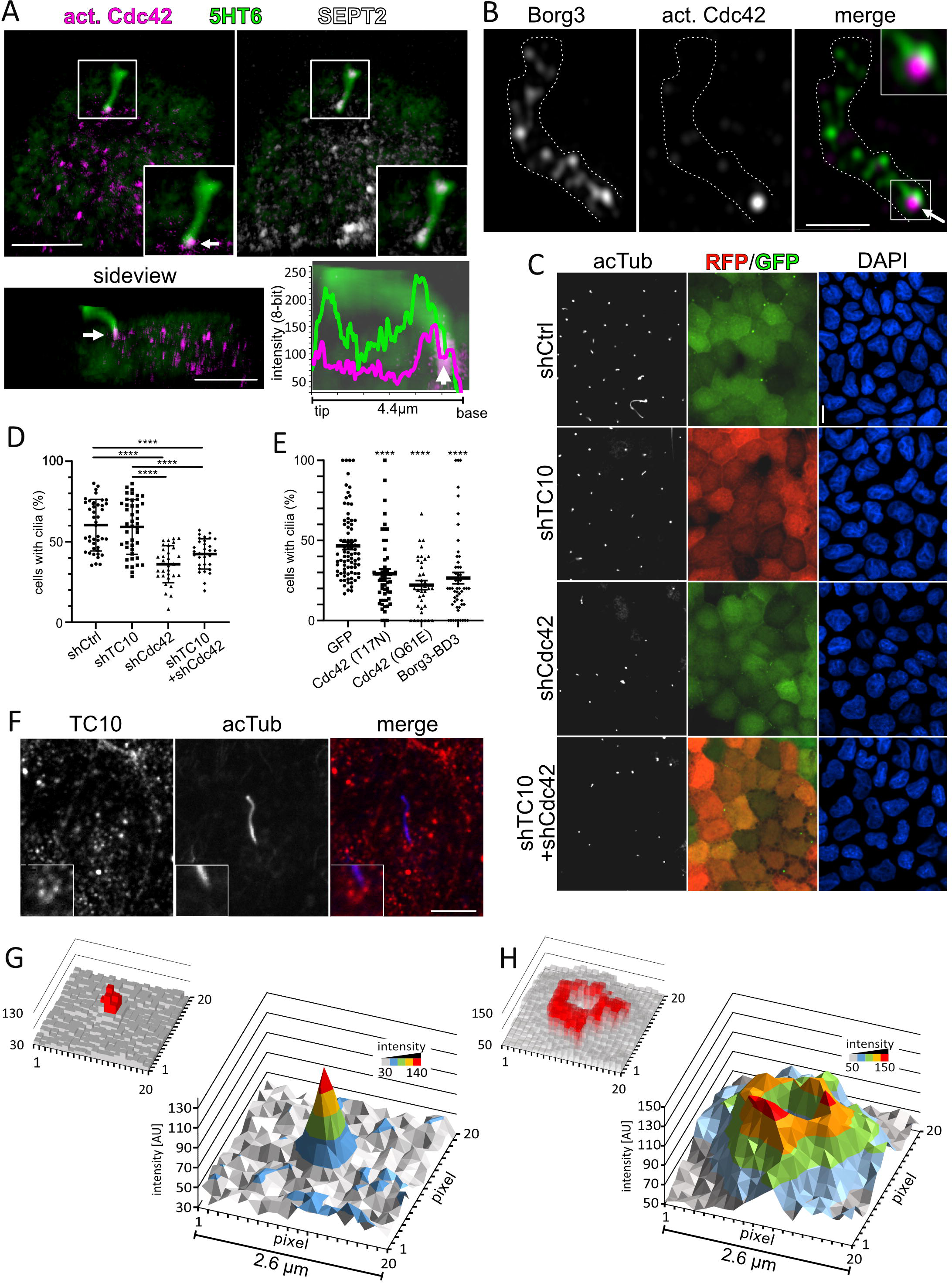
Formation of cilia is regulated by Cdc42. **A)** Representative confocal image of MDCK cells transfected with 5HT6-tomato to visualize the primary cilium immunostained for SEPT2 and active Cdc42. The images show the top view on the cilium (upper row), the side view (bottom left) and a 3D-linescan. Arrows in the images and the linescan indicate accumulation of active Cdc42 at the base of the cilium. Scale bar 5 μm. **B)** Representative SIM image of IMCD cells immunostained for endogenous Borg3 and active Cdc42. Arrow indicates accumulation of active Cdc42 with surrounding Borg3 at the ciliary base. Borg3 also locates along the axoneme. Scale bar 1 µm. **C)** Representative confocal images of stable MDCK KD cell lines immunostained for acetylated tubulin. Cells that express shRNA targeting Cdc42 and/or TC10 or non-silencing control (ns control) are marked by GFP and/or RFP fluorescence. Cells were serum starved for 48h before fixation. Nuclei were stained with DAPI. Scale bar 5 μm. **D)** Graph shows quantification of cilia formation in TC10 and Cdc42 KD cells (representative images shown in C). Data are given ±SEM, ≥30 fields of view were analyzed n=3. **E)** MDCK cells were transfected with Cdc42 (T17N)-GFP, Cdc42 (Q61E)-GFP, the septin binding domain BD3 or GFP as control and were starved for 48h. Cells were stained for acetylated tubulin and nuclei. Data are given ±SEM, >30 fields of view were analyzed, n=4. **F)** Representative confocal images of IMCD cells immunostained for TC10 and acetylated tubulin. Cells were serum starved 48h before fixation. Scale bar 5 μm. **G)** 3D-intensity surface-blot shows quantification of active Cdc42 immunofluorescence intensity at the ciliary base in a 2.6 μm square (400 pixels). 15 images of active Cdc42 at the base of individual cilia were analyzed. For all cilia the z-plane sectioning the ciliary base was used. All images were centered on the base of the cilium. The fluorescence intensity of each pixel was measured for each image and summed up to generate a 3D-graph (AU = arbitrary units). The Anova pixel analysis (small graph, left) shows every pixel in red with statistically elevated intensity. n=15. **H)** Same quantification as in G) for TC10 intensity at the cilium. Here a TC10 antibody was used that recognizes active- and inactive TC10. n=15.

To further test the paradigm that Cdc42 is important for ciliogenesis we used stably expressed shRNAs targeting Cdc42 in MDCK cells. The KD of Cdc42 reduced the percentage of ciliated cells up to 45% compared to the control (Figure 4 D and Figure S3 D). Next, we wanted to assess the impact of the nucleotide exchange within Cdc42 on the process of cilia formation. We hypothesized that switching Cdc42 between the active GTP-bound form and the inactive GDP-bound form will influence the efficacy of Borg3 recruitment and ciliogenesis. MDCK cells were transfected with GFP-tagged constitutively active (Q61E) or inactive (T17N) mutants of Cdc42 and the number of transfected cells with cilia was quantified (Figure 4 E). The expression of dominant negative Cdc42 (T17N) and dominant active Cdc42 (Q61E), both resulted in a reduced number of ciliated cells. Moreover, expression of the septin binding domain of Borg3 (BD3), that lacks the regulatory Cdc42 interaction site, led to a reduction in ciliogenesis to a similar extent (50%). We reason that the tight spatio-temporal regulation of Cdc42 and its downstream effector Borg are important for septin regulation during ciliogenesis. A global cellular deregulation of Cdc42 has a negative influence on the formation of cilia.

Besides Cdc42, Borg proteins are also interacting with TC10 known as RhoQ (23). This small Rho GTPase is a close homolog of Cdc42. Borg3, however, is exclusively regulated by Cdc42 (23). In several different cellular functions TC10 and Cdc42 have been shown to be redundant (34, 35). TC10 can be found predominantly at the cell membrane and vesicles where it is essential in regulating exocytosis in adipocytes and neurons (36, 37). However, the function of TC10 in Borg regulation and downstream septin regulation at cilia was not studied yet. Immunofluorescence microscopy and expression of GFP-TC10 revealed that TC10 accumulates around the base of cilia (Figure 4 F and S3 E). A quantitative spatial analysis of active Cdc42 (Figure 4 G) and TC10 (Figure 4 H) fluorescence at the base of cilia revealed a significant difference in the localization preference of the two Rho GTPases. Whereas active Cdc42 can be found predominantly in a spot at the base of cilia, TC10 spares that area and surrounds the base of cilia in small clusters. We assume that TC10 positive vesicles arrange around the base or transition zone of the cilium. Notably, Figure 4 H does not imply that TC10 forms a solid ring around cilia, it shows the area where most of TC10 fluorescence occurred in the optical plane of the transition zone as a sum from 15 cilia. These results suggest that TC10 and Cdc42 might have different functions during ciliogenesis. Moreover, the non-overlapping localization of TC10 and Borgs suggests that TC10 is not involved in the localization of Borg3 to cilia. Similar to Cdc42 independent of its activity state (S3 B and C), we observed TC10 within the cilium, as well. However, this was restricted to smaller clusters (Figure S4 F).

To further analyze the role of the RhoGTPases in ciliogenesis upstream of Borg3, stable KDs of TC10, Cdc42, and both together were generated in MDCK cells (Figure S4 D) Single KD of TC10 had no effect on the number of ciliated cells. The double KD of TC10 and Cdc42 reduced ciliogenesis in the same way as the single KD of Cdc42 (Figure 4 C and D), so there was no additional effect. These data further indicate that Cdc42 and not TC10 is the upstream regulator of Borg3 localization at cilia. It is tempting to speculate that Cdc42 and Borg3 affect the formation of cilia by the regulation of septin recruitment and possibly septin assembly. Since Borg3 interacts with the active form of Cdc42 (23), we hypothesized that the switching of Cdc42 between the inactive and active state is necessary to recruit Borg3 in sufficient amounts to fuel septin accumulation and subsequent assembly at cilia.

### Borg3 regulates Septin dynamics at cilia

We analyzed the recruitment of septins to cilia of Borg3 KO cells to observe changes in the absence of this septin binding protein (29) (Figure 5 A-D). SEPT2 (Figure 5 A and B) or SEPT7 (Figure 5 C and D) were stained by immunofluorescence in WT and Borg3 KO cells. An acetylated tubulin staining was used as a mask to evaluate septin fluorescence intensity in the ciliary compartment. The intensity of SEPT2 and SEPT7 were significantly reduced in Borg3 KO cells. These data suggest that Borg3 is involved in the recruitment of septins to the ciliary compartment or in the regulation of septin assembly in cilia.

**Figure 5.**
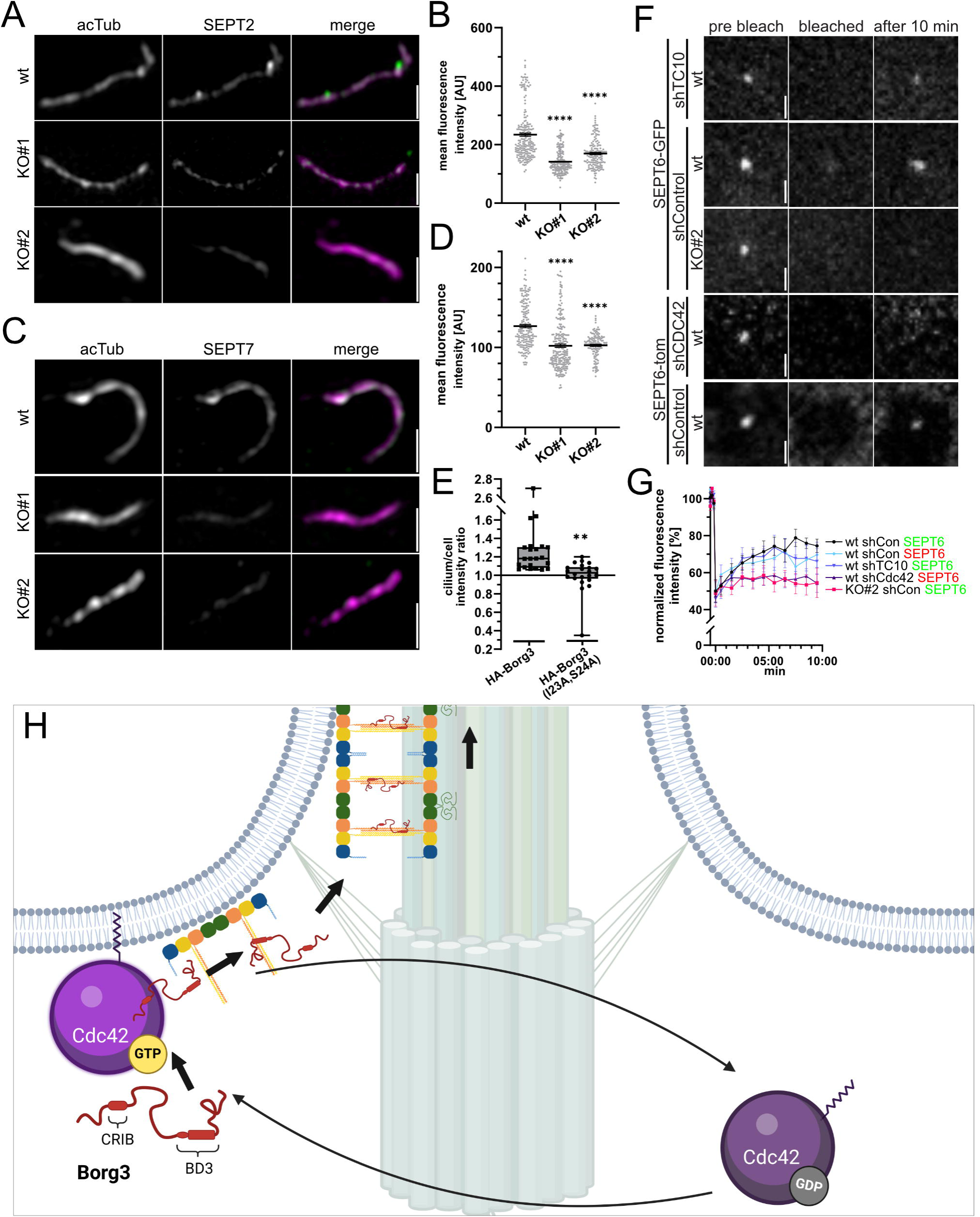
Localization Septins and Borg3 to the primary cilium depends on Cdc42. **A)** Representative SIM images of cilia of IMCD wt and Borg3 KO cell lines immunostained for endogenous SEPT2 and acetylated tubulin. Cells were serum starved 48h before fixation. Scale bar 1 µm **B)** Quantification of SEPT2 immunofluorescence average intensity in cilia from IMCD cells treated as in A. Acetylated tubulin was used as mask for intensity measurements (AU = arbitrary units). Data are given ±SEM. ≥9 random fields of view were analyzed from n=3. **C)** Representative SIM images of cilia of IMCD wt and Borg3 KO cell lines immunostained for endogenous SEPT7 and acetylated tubulin. Cells were serum starved 48h before fixation. Scale bar 1 µm **D)** Quantification of SEPT7 immunofluorescence average intensity in cilia from IMCD cells treated as in C. Acetylated tubulin was used as mask for intensity measurements (AU = arbitrary units). Data are given ±SEM. ≥9 random fields of view were analyzed from n=3. **E)** 3xHA-Borg3 and 3xHA-Borg3 (I23A, S24A) were expressed in IMCD cells starved for 48h. Cells were immunostained for the HA-tag and for Arl13b. Confocal images were acquired. A mask was generated by the Arl13b channel to measure the HA average fluorescence intensity in the cilium. This value was divided by the average intensity adjacent to the cilium in the focal plane to yield a cilium/cell intensity ratio. Data are given ±SEM. n=4. **F)** Representative confocal images of IMCD cell lines stably expressing shRNA targeting Cdc42, TC10 or a non-targeting shRNA control as well as the IMCD Borg3 KO#2 cell line stably expressing the non-targeting shRNA control transfected with SEPT6-GFP (for shTC10 and respective control) or SEPT6-tom (for shCdc42 and respective control) 24h before imaging. SiR-tubulin was added to the cells as a ciliary marker 1h before imaging. Cells were bleached with the corresponding laser until 50% of the previously observed fluorescence intensity inside the cilium was lost and recovery was observed for 10min. **G)** Quantification of ciliary SEPT6 intensity in IMCD wt and Borg3 KO#2 cell lines treated as in F. Data are given ±SEM. n=10. **H)** Scheme of proposed mechanism of the Cdc42 and Borg3 mediated entry of septins to the primary cilium. Cdc42 constantly cycles from inactive GDP bound to active GTP bound state at the ciliary base. Active Cdc42 is recruited to the membrane by its CaaX motif modified by isoprenylation. Borg3 binds to activated Cdc42 by its CRIB domain and is thereby enriched in the proximity of septin octamers at the membrane. Borg3 interacts with the c-terminal tails of SEPT6 and 7 and aids in formation of septin filaments and subsequently more complex structures enrich in the ciliary compartment. The illustration was created with BioRender.

To further analyze the function of Borg3 on septin assembly at cilia a Borg3 variant was used that is not able to interact with Cdc42 by I23A,S24A mutations in the CRIB-domain (23) (Figure 5 E and Figure S4 G). After expression of HA-tagged versions of wt and mutant Borg3, significantly less mutant Borg3 could be detected in the ciliary compartment compared to the cytosol.

To further analyze the impact of Borg3 on septin dynamics at cilia, we performed bleaching experiments (Figure 5 F and G). SEPT6-GFP was expressed in wt and KO#2 cells stained with SiR-tubulin to identify cilia. We bleached shorter cilia because we aimed to bleach the complete cilium and monitor SEPT6 entry from the cytosolic pool without excessively reducing fluorescent SEPT6 in the cytosol. Smaller cilia might have stronger SEPT dynamics due to growth phases. Bleaching of SEPT6 revealed a more efficient entry of SEPT6-GFP in the ciliary compartment when functional Borg3 was present. Notably, the entry of SEPT6-GFP was rather slow as it took almost 10 min to reach 80% of the pre-bleach fluorescence intensity. These experiments suggest that Borg3 plays a role in septin dynamics at cilia. Borg3 is regulated or recruited by Cdc42 that has to cycle between the inactive and active state. Bleaching cells expressing shRNAs targeting Cdc42 or TC10 (Figure S4 H) further supported the theory that Cdc42 is the upstream regulator of septin localization to primary cilia. Cells with Cdc42 KD showed reduced fluorescence recovery compared to the respective control mimicking the effect of Borg3 KO. TC10 KD had no effect on fluorescence recovery (Figure 5 F and G) compared to the respective control.

## Discussion

Here, we identify Borg3 as a new regulator of primary cilia formation. We show that interference with Borg3 function by either KD or KO reduces the formation of primary cilia. Borg3 can be found at the base and within the cilium. It colocalizes with septin structures in these areas. Over the years several ciliary proteomes were analyzed. Septin family members were regularly identified, while Borg proteins were never a major hit in these studies. Borg3 might be a difficult target for mass spectrometry since tryptic digests results in a very long peptide and another peptide that is prone to crosslinking. Thus, only a low sequence coverage can be expected in mass spectrometry.

Only two domains of Borgs have an assigned function. The first functional domain is a CRIB-domain which mediates interaction with Cdc42. However, Borg3 seems to be no classical Cdc42 effector, as it might not exert an enzymatic function upon activation. Instead, active Cdc42 might be solely necessary to localize Borg3 (11, 14, 25).

The second functional domain of Borg3 is the BD3 domain. Previous studies demonstrated that the BD3 domain interacts with the C-terminal domains of SEPT6 and SEPT7 (22, 29). This interaction may not be restricted to a single filament it may also include adjacent filaments which could promote the formation of bundles and higher order structures. Moreover, it was shown that the BD3-domain interaction with the c-terminal domain of SEPT6 and SEPT7 can induce polymerization of septin oligomers into filaments in vitro (29).

In line with this notion, we identified Cdc42 as a regulator of ciliary septin assembly and Borg3 recruitment. Locking Cdc42 in an active or inactive state as well as downregulation by KD reduced cilia formation and recruitment of Borg3 and septins. The spatio-temporally regulated switch of Cdc42 between the inactive state and the active state appears essential for the Borg3 assisted septin recruitment or assembly. Moreover, a repetitive cycling between the inactive and active state could amplify the process of Borg3 attraction by constantly recruiting and releasing. Thereby, a concentration of Borg3 and septins could be achieved that favors septin assembly at cilia. To this end, a Cdc42 guanine nucleotide exchange factor and GTPase-activating protein (GAP) are necessary at cilia. Tuba and intersectin 2 are two promising candidates as GEFs (32, 33). Moreover, GAPs for Rho-GTPases have appeared in ciliary proteomes (38, 39) and might be potential interaction partners for septins (40).

Most in vitro studies use a shift in ionic strength from high to low for induction of septin polymerization (41), which is probably not or not the only physiological key to control septin dynamics in cells. Notably, in the presence of the BD3 domain, septin polymerization is independent of a shift in ionic strength in vitro (29). To this end, we propose a mechanism by which a GTPase-dependent switch at the base of cilia increases Borg3 concentration to trigger efficient septin assembly at cilia (Figure 5 H). This raises the question how a selectivity for Borg3 is generated as it was demonstrated that all Borgs share the two conserved functional CRIB- and BD3 domains. We observed that active Cdc42 is enriched at the base of the cilium. This confined spot is surrounded by an area with higher TC10 activity. As it was shown early, Borg3 is the only Borg protein that is exclusively regulated by Cdc42 and not by TC10 as well (23). It is therefore tempting to speculate that the other Borgs are also recruited to the periphery of the cilium while Borg3 is exclusively attracted directly to the cilium. This does not rule out that TC10, other Borgs and downstream septins have alternative cilia-related functions, for example during vesicle tethering and subsequent exocytosis (42, 43) which could play an important role in directed transport of membrane proteins to the cilium.

The localization of active Cdc42 at the base of cilia with subsequent Borg3 recruitment could provoke additional synergistic effects with the adjacent membrane (Figure 5 H). An isoprene derivative is linked to the cysteine residue of the C-terminal CaaX-box of Cdc42 which anchors the Rho-GTPase to the membrane (44). Septins have a preference to accumulate at membranes and in particular with curved membranes (45, 46) found at the base of cellular protrusions. Thus, additional means of enrichment are available that can reinforce the process by concentrating all necessary components at the site of polymerization. Notably, in fungi Cdc42 at the plasma membrane and its upstream regulators define the site of septin recruitment and assembly during bud formation (47).

Taken together, the cytoskeletal partners of septins, actin and microtubules have a large number of binding proteins that facilitate and fine-tune their polymerization. For example, formins and +end binding proteins that regulate actin- and microtubule polymerization, respectively. It is plausible that the septin cytoskeleton employs similar mechanisms of regulation. Thus, Borg proteins are currently the most documented candidates for such functions for the septin cytoskeleton. We report that Borg3 takes a vital role in the regulation of septin accumulation at cilia. Notably, further studies are necessary to analyze the higher order structures septins can form within cilia and how Borgs influence these structures. Moreover, further studies are necessary to elucidate the involvement of Borgs in the other cellular functions attributed to septins. The exact molecular mechanism how Borg proteins facilitate septin filament- and higher order structure assembly needs further investigation to advance our knowledge of these septin binding proteins.

## Methods

### Cell Culture and Transient Transfections

MDCK cells were cultured in Modified Eagle’s Medium (MEM) supplemented with 10% FCS, 1% penicillin/streptomycin and 1% nonessential amino acids. Platinum-E cells and IMCD cells were cultured in DMEM/F12 supplemented with 10% FCS, 1% penicillin/streptomycin. 1% nonessential amino acids and 1% sodium pyruvate (Biochrom) were additionally added for Platinum-E cells. Mycoplasma contamination was excluded by monthly testing.

To enhance ciliogenesis cells were placed in their respective growth medium without FCS for 48h. For immunostainings, cells were plated on HCl-washed coverslips or on µ-Slide 8 wells (Ibidi). For live-cell imaging, cells were plated on glass-bottom dishes (Greiner).

Cells were transfected using Lipofectamine 2000 (Invitrogen) according to the manufacturer’s protocol. When necessary, transfected cells were enriched by fluorescence activated cell sorting with a BD Melody with 488 nm and 561 nm laser lines.

### shRNAs

The shRNAs for SEPT2, for SEPT7 and for SEPT9 were cloned into the pSUPER.retro.neo+GFP vector (Oligoengine).

The Cdc42 shRNA were inserted into the pGIPZ lentiviral vector with GFP (Clone-Id: V3LHS_641566 GE Dharmacon and Clone-Id: V2LHS_261933 GE Dharmacon). TC10 shRNA was inserted into pSMART lentiviral vector with Turbo RFP. Following non-silencing control was used: pGIPZ non-silencing lentiviral shRNA control (# RHS4348 GE Dharmacon). Sequences are shown in Table 1.

### Lentiviral Transfection

Packaging of the lentiviral vectors pGIPZ-GFP shRNA Cdc42 #2, pGIPZ-GFP non-silencing control, pSMART-TurboRFP shRNA TC10 was performed using transient transfection into Platinum-E cells (a gift from Prof. Toshio Kitamura, University Tokyo, (48)) together with five packing plasmids: pTLA1-Pak, pTLA1-Enz, pTLA1-Env, pTLA1-Rev and pTLA1-TOFF (Thermo-Fisher). 1.5 x 10^6^ Platinum-E cells were plated onto 10 cm dishes and grown for one day. Culture supernatants of the lentivirus producing cells were collected. The medium from sub-confluent MDCK cells was removed and replaced by a 1:1 mixture of lentivirus containing medium and growth medium, supplemented with 8 µg/ml polybrene. After 48 hours selection of transduced cells was initialized by 3 µg/ml Puromycin. The infection efficiency of the lentivirus was determined by green or red fluorescent protein (GFP and RFP) expression. Double KD cells were first transduced with pGIPZ-GFP shRNA Cdc42#2. The expression of each construct was detected using a fluorescence microscope.

### CRISPR/Cas9 mediated Knockout of Borg3

Borg3 KO cell lines were generated according to Ran et al. (27). In brief, targeting sequences (Table 1) for the guide RNAs were cloned into pSpCas9(BB)-2A-GFP (PX458)-plasmid (from addgene, #48138). 500 ng of the plasmid were introduced to 90% confluent IMCD cells. Cells were sorted and seeded as single cells. New cell lines carrying the desired mutation were identified by sequencing. Nucleotide sequences are shown in Table 1.

### Antibodies, Fluorescent Dyes, and Fluorescent Proteins

The used antibodies and dyes are listed in Table 1.

dsRed-cent2 (Plasmid #29523) (49) and GFP-TC10 (Plasmid #23232) (50), pEGFPN3-5ht6 (#35624) (51), L13-Arl13bGFP (#40879) (52), were obtained from addgene for direct use or further subcloning. 3xHA-Borg3 was provided by Ian Macara (23). Other plasmids were previously used in (12, 24)

### Immunostaining

Cells were washed with PBS, fixed for 15 min with 4% formaldehyde in PBS, permeabilized (10min) with 0.15% Triton X-100 in PBS, and blocked by 1% BSA or normal goat serum in PBS for 30min. Cells were incubated with the primary antibody for 60-90min at RT. Cells were washed with PBS and incubated with the suitable secondary antibody for 1h. Cells on coverslips were washed with in PBS, water, dried, and embedded with Prolong Diamond (Thermo Fisher Scientific). Cells in µ-slide 8 wells were washed and left in PBS. µ-slides were used whenever reliable identification of the ciliary base was necessary, since a large fraction of cilia stay upright. For nuclear staining DAPI (Sigma) was added to the normal staining or embedding medium supplemented with DAPI (Thermo Fisher Scientific) was used. For parallel immunostainings of Borg3 and different septins (all relevant antibodies produced in rabbit), the FlexAble kit for primary antibody labelling was used according to manufacturer’s protocol. Critical stainings were verified by a staining according to (53) with PFA in Cytoskeleton buffer.

### Imaging

Cells were analyzed with an Axio Observer microscope (Carl Zeiss), driven by Visiview (Visitron) imaging software with plan-apochromat objectives, a Yokogawa CSU-X1 spinning disk confocal head with emission filter wheel, 405-, 488-, 561-,640 nm laser lines, and a Coolsnap HQ II digital CCD camera or prime bsi sCMOS camera (Teledyne Photometrics). Alternatively, cells were imaged with a LSM800 confocal microscope (Carl Zeiss), multiplex airyscan / GaAsP-detectors and 405-, 488-, 561-,640 nm laser lines or an Elyra 7 structured illumination super resolution microscope with dual PCO edge sCMOS cameras. For live-cell imaging, cells were incubated in a chamber with humidified atmosphere (5.5% CO_2_) at 37°C on the microscopes mentioned above.

For real-time quantification of cilia formation, a Lionheart FX automated microscope (Agilent BioTek) with environmental control was used.

### Image processing and analyis

SIM images were processed with ZEN Black software. The theoretical optical transfer function given by the manufacturer was used. ‘Baseline shift’ was deactivated for all reconstructed images as a standard, and thus the threshold was set manually into the first peak appearing in the histogram for representative images. No grey values of the histogram were influenced or cut off for further quantitative analysis. Maximum-resolution enhancement by minimal artefact structure (‘hammer finish artefacts’) was targeted by comparing background patterns with structure signals for representative images.

All microscopy images were further processed with Imaris or Metamorph software.

To quantify formation of cilia in fixed cells, cilia and nuclei of all cells per field of view were counted. Nuclei were stained with DAPI and cilia were stained with acetylated tubulin as well as Arl13b. The number of cilia in relation to the total number of nuclei was evaluated. For transfected cells, only the total number of transfected cells with cilia were counted.

For intensity quantifications of immunofluorescence stainings a 2D- or 3D mask was generated by a ciliary marker (i.e. Arl13b or acetylated tubulin). The mask was applied to the channel of interest to exclusively measure fluorescence intensities at the cilium. The mean fluorescence intensity for each primary cilium was used for subsequent analyses.

For quantitative mapping of fluorescence intensities at the base of cilia, a 20x20 pixel square was centered at the base of the cilium. Intensities of all 400 pixel were measured and summed up for each pixel coordinate of the 15 cilia analyzed. All pixel values were compared by one way ANOVA with Tukeýs posttest to identify statistically significant elevations in fluorescence intensity in the averaged image. The generated 3D-blot for each pixel gives an overview where fluorescence signal accumulated.

In order to identify formed cilia in real time long-term video microscopy a fluorescence intensity- and size-dependent filter was used to discriminate cell debris from newly formed cilia. Only intensities following the first peak in the histogram were evaluated and subsequently only counted if they covered an area of 4-90 pixels in size.

### Fluorescence Recovery after Photobleaching

IMCD cell lines containing shRNAs or a non-targeting control were transfected with SEPT6-GFP or SEPT6-tom as to not interfere with the fluorescence needed to identify shRNA expressing cells. Primary cilia were identified by SiR-tubulin and an area of 2 µm by 2 µm centered on the primary cilium was bleached with the corresponding laser until 50% of the fluorescence intensity was lost. Recovery was observed for 10 min inside the primary cilium using the SiR-tubulin fluorescence as positioning and size marker. During imaging an automated focus was used. Bleaching was performed using the corresponding laser for the transfected protein (560 nm for SEPT6-tom and 488 nm for SEPT6-GFP) with 10-15% laser intensity as needed from the fluorescence intensity observed before bleaching. Bleaching was stopped automatically once 50% of initial intensity was bleached.

### Statistics

Student’s *t* test was applied when two groups with normal distribution had to be compared. Whenever more than 2 groups were compared a Oneway ANOVA with Tukey posttest was used. Statistical evaluation was performed with GraphPad Prism. *P* values <0.05 were considered statistically significant and marked with an asterisk (\**P* < 0.05; \*\**P* < 0.01; \*\*\**P* < 0.005; \*\*\*\**P* < 0.001).

## Acknowledgements

We thank Feng Zhang, Joseph Gleeson, Channing Der, Kirk Mykytyn and Tamara Caspary for Addgene plasmids. We thank Ian Macara for 3xHA-Borg3 plasmid. Work was supported by Deutsche Forschungsgemeinschaft (DFG) with a research grant (SCHW 1708/2-1) to C.S.. We thank Otilia Wunderlich and Peter Gebhardt for technical assistance.

## Data availability statement

The data that support the findings of this study are available from the corresponding author upon request.

