## Supplementary Information for "Borg3 controls septin recruitment for primary cilia formation"

### Supporting Information

#### Legends

**Table 1** Table containing used antibodies

**Table 2** Table containing used oligonucleotides

**Fig. S1 A, B)** Representative SIM image of IMCD cells immunostained for endogenous SEPT2 and SEPT9, A) and B) respectively, and acetylated tubulin. Septins are localized at the transition zone of primary cilia (arrow). **C)** As in A) and B) but with immunostaining for SEPT7 forming a partial transition zone ring (arrow). **D)** MDCK cells were transfected with shRNA targeting SEPT9. KD was confirmed by western blot. GAPDH was used as a loading control (35.9 kDa). **E)** Representative SIM images of MDCK cells treated as in D and immunostained for endogenous SEPT2 and acetylated tubulin. Cells were serum starved 48h before fixation. Scale bar 1 µm. **F)** MDCK cells were transfected as in D. Cells were starved for 48h, fixed and stained for acetylated tubulin by immunofluorescence. GFP additionally expressed from the shRNA vector was used to identify KD cells. The percentage of cells with a cilium was quantified. The average of the controls was set to 100% and used for normalization. Data are given as mean ±SEM. ≥20 random fields of view were analyzed from n=3. **G, H)** MDCK cells transfected with shRNA targeting SEPT7 G) and SEPT2 H) respectively. KD was confirmed by western blot. GAPDH was used as a loading control (35.9 kDa). **I)** IMCD cells transfected with siRNA targeting Borg1, Borg2 or Borg3 48 or 72h before harvesting. Efficiency of KD was verified by quantitative real time PCR. IMCD cell transfected with a non-targeting siRNA 48h before harvesting were used as control. n=3. **J)** Representative confocal images of subconfluent MDCK cells transfected with GFP-Borg1-, 2- or, 3 and immunostained for endogenous SEPT2 and α-tubulin. Septin filaments colocalize with all three expressed Borg proteins. Scale bars 10 µm.

**Fig. S2 A)** Sequencing results of the generated IMCD Borg3 KO cell lines (KO#1 and KO#2) as well as wildtype IMCD cells (wt). In KO#1 a heterozygous mutation results in ambiguity from the mutation site. Using the wt sequence the sequences for the individual allels are identified and written above. Newly introduced nucleotides (compared to the wt sequence) are indicated by underlining, loss of nucleotides is indicated by arrows and displaying of lost nucleotides. KO#2 homozygous mutation is identified by comparison with the wt sequence. Newly introduced nucleotide is indicated by underlining. **B)** Representative confocal images of IMCD Borg3 KO cell line KO#1 immunostained for endogenous Borg3 and acetylated tubulin. KO verification by immunofluorescence staining. **C)** Representative confocal images of the computer aided identification of primary cilia used to obtain Fig. 3 C and D. Images were gated for high intensity fluorescent particles followed by a count of particles with distinct size requirements (area: 4-90 pixels).

**Fig. S3 A)** Representative confocal image of MDCK cells transfected with GFP-Tuba and DsRed-Centrin2. Cells were serum starved 48h before fixation. Scale bar 10 µm. **B)** Representative confocal image of MDCK cells transfected with GFP-Cdc42 and immunostained for SEPT2 and acetylated tubulin. Scale bar 10 µm. **C)** Representative SIM image of primary cilia in IMCD cells immunostained for endogenous Cdc42 and Arl13b. Scale bar 1 µm. White dashed line indicates intensity line scan below. **D)** Western blot verification of MDCK cell lines stably expressing shRNA targeting TC10 and/or Cdc42, respectively. GAPDH (35.9 kDa) was used as loading control. **E)** Immunofluorescence of IMCD cells transfected with GFP-TC10 and stained for SEPT2 and acetylated tubulin. The images for SEPT2 and acetylated tubulin display a focal plane at the axoneme of the cilium. The image of TC10 accumulation is at the apical cell membrane of the same z-stack. Scale bar 5 µm. **F)** Representative SIM image of primary cilia in IMCD cells immunostained for endogenous TC10 and acetylated tubulin. Scale bar 1 µm. **G)** Representative confocal images of IMCD cells transfected with 3xHA-Borg3 and 3xHA-Borg3 (I23A, S24A) and immunostained for the HA-tag and endogenous Arl13b. Scale bar 5 µm**. H)** Western blot verification of IMCD cell lines stably expressing shRNA targeting Cdc42 or TC10, respectively. α-tubulin and GAPDH (35.9 kDa) were used as loading controls, respectivley.

**Movie 1** IMCD cells fixed in µ-slides (Ibidi) and stained for Borg3 (green) and acetylated tubulin (red) by immunofluorescence. Z-stack was 3D-rendered and animated by imaris. Scale bar indicates magnification. Stills of zoomed in cilium in Fig. 2 D.

**Movie 2** IMCD cell stained for Borg3 (green) and acetylated tubulin (red) by immunofluorescence. Z-stack was 3D-rendered and animated by imaris. Scale bar indicates magnification.

**Table 1**

| **Antibodies** | | | | |
|---|---|---|---|---|
| Antibody | Catalog no. | clone | used for | supplier |
| CDC42EP1 | HPA006379 | | IF, WB | Sigma-Aldrich |
| CDC42EP2 | HPA038562 | | IF, WB | Sigma-Aldrich |
| CDC42EP3 | HPA061792 | | IF, WB | Sigma-Aldrich |
| CDC42EP5 | PA5-106637 | | WB | ThermoFisher |
| CDC42EP5 | HPA043449 | | IF | Sigma-Aldrich |
| SEPT2 | HPA018481 | | IF | Sigma-Aldrich |
| SEPT7 | HPA029524 | | IF | Sigma-Aldrich |
| SEPT6 | sc-20180 | H-60 | IF | Santa Cruz |
| SEPT9 | HPA042564 | | IF | Sigma-Aldrich |
| Rab8 | 865401 | 18073A | IF | BioLegend |
| CEP164 | sc-515403 | E-9 | IF | Santa Cruz |
| Sec8 | MABC570 | 2E12 | IF | Merck Millipore |
| acetylated α Tubulin | sc-23950 | 6-11B-1 | IF | Santa Cruz |
| CDC42 | sc-8401 | B-8 | IF | Santa Cruz |
| active CDC42 | 26905 | | IF | New East Biosciences |
| TC10 | T8950 | | IF, WB | Sigma-Aldrich |

**Table 2**

| internal name | sequence | used for | catalog no. |
|---|---|---|---|
| mCdc42ep2_F | TCCCCATCTATTTGAAACGTGG | qPCR | |
| mCdc42ep2_R | CCGCTGTTCCTGGAAGGAG | qPCR | |
| mCdc42ep3_F | CCAAGACCCCAATTTACCTGAAA | qPCR | |
| mCdc42ep3_R | CCCTCTTTGCCGATGTGTATAGT | qPCR | |
| mCdc42ep5_F | GGGATGCCCACCCTAGAGT | qPCR | |
| mCdc42ep5_R | TGGAGGTCAGCATTTGAGCAG | qPCR | |
| Borg3 p3 fw | CACCGAGCCGCTTCTTGGGTTGTGC | CRISPR/Cas9 | |
| Borg3 p3 rv | AAACGCACAACCCAAGAAGCGGCTC | CRISPR/Cas9 | |
| Borg3 p4 fw | CACCGTGCACAACCCAAGAAGCGGC | CRISPR/Cas9 | |
| Borg3 p4 rv | AAACGCCGCTTCTTGGGTTGTGCAC | CRISPR/Cas9 | |
| Borg3 p5 fw | CACCGCCCTGCACAACCCAAGAAG | CRISPR/Cas9 | |
| Borg3 p5 rv | AAACCTTCTTGGGTTGTGCAGGGC | CRISPR/Cas9 | |
| Borg3 seq. fw | gatcaggtacagttatgggcg | sequencing | |
| Borg3 seq. rv | GTGGAGTGCTGGGAGGGAG | sequencing | |
| Borg3 seq. 2 fw | CAGGAGGGACCTTGAGAACCT | sequencing | |
| Borg3 seq. 2 rv | CGACTCAGGAATGAGGTGTCC | sequencing | |
| AllStars Neg. Control siRNA | | Knock Down | 1027292 |
| Cdc42ep2#1 | GACCUUCCCUUCCAGUUUA | Knock Down | D-044823-01 |
| Cdc42ep2#2 | GAUUAUGGAUCACGACCUA | Knock Down | D-044823-02 |
| Cdc42ep2#3 | UGGCGGAGAUGACAUGUUU | Knock Down | D-044823-03 |
| Cdc42ep2#4 | CGUGCAGAUUCCUACAUA | Knock Down | D-044823-04 |
| Cdc42ep3#1 | GGAGCAAAGUAGUCUAUUA | Knock Down | D-046421-01 |
| Cdc42ep3#3 | GAUCUUGGGCCUUCACUUU | Knock Down | D-046421-03 |
| Cdc42ep5#1 | GGAGCACUCUCGAUCUCAG | Knock Down | D-063228-01 |
| Cdc42ep5#3 | CGACGUCACGGGUCUGUAG | Knock Down | D-063228-03 |
| shSEPT2 | GGAGAACATCGTGCCCGTC | Knock Down | |
| shSEPT7 | CACAGTATCCTTGGGGTGT | Knock Down | |
| shSEPT9 | GTCCATCACGCACGATATT | Knock Down | |

| | | | |
|---|---|---|---|
| shScramble | GATCTGATCGACACTGTAA | Knock Down | |
| shCdc42 #1 | CAGGAGACATGTTTTACCA | Knock Down | V3LHS_641566 |
| shCdc42 #2 | TCTGTCATAATCCTCTTGC | Knock Down | V2LHS_261933 |
| shTC10 | CACGTAATCAAACAACAGT | Knock Down | V3SVMM08_11985849 |
| pGIPZ non-silencing lentiviral shRNA control | | Knock Down | RHS4348 GE |

**Fig. S1**

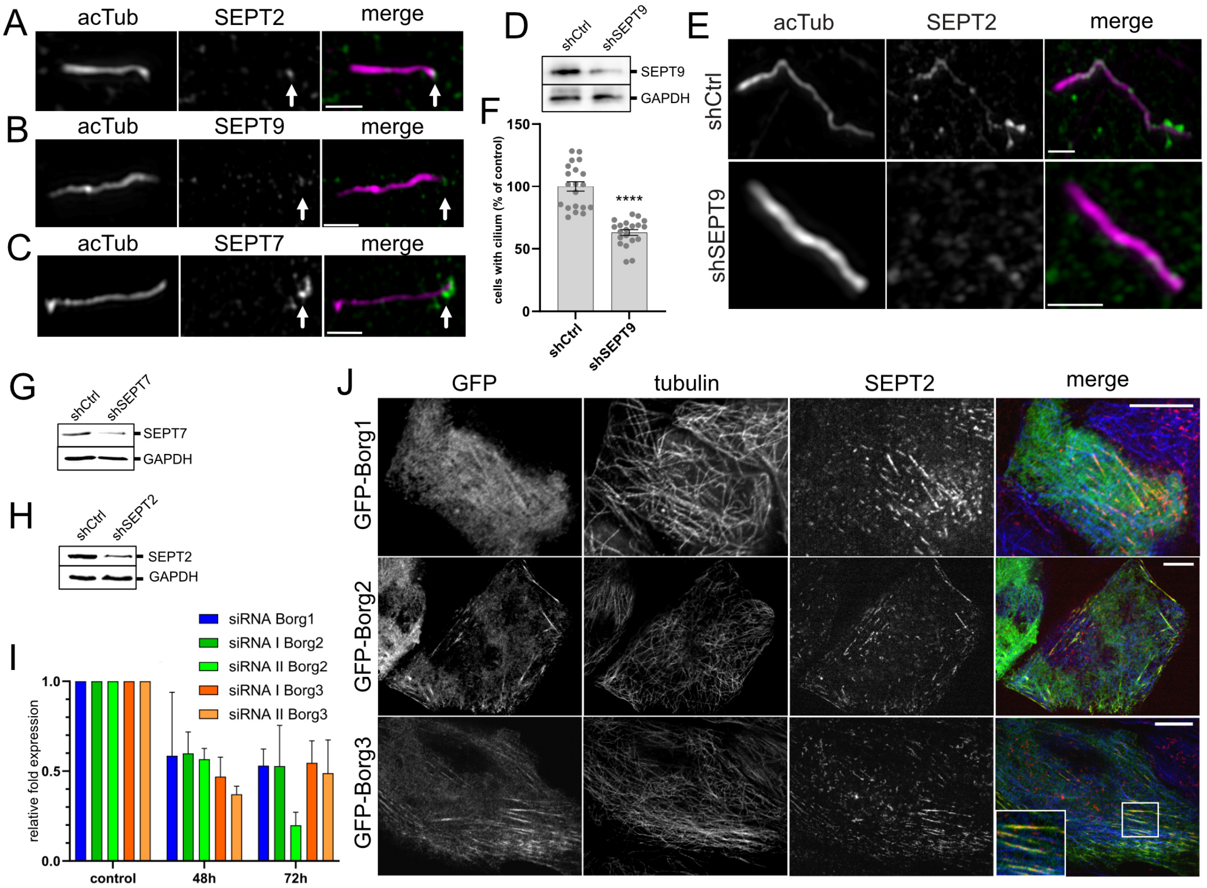
A
acTub
SEPT2
merge
B
acTub
SEPT9
merge
C
acTub
SEPT7
merge
D
shCtrl
shSEPT9
SEPT9
GAPDH
F
cells with cilium (% of control)
150
100
50
0
****
shCtrl
shSEPT9
E
acTub
SEPT2
merge
shCtrl
shSEPT9
G
shCtrl
shSEPT7
SEPT7
GAPDH
H
shCtrl
shSEPT2
SEPT2
GAPDH
I
siRNA Borg1
siRNA I Borg2
siRNA II Borg2
siRNA I Borg3
siRNA II Borg3
relative fold expression
1.0
0.5
0.0
control
48h
72h
J
GFP
tubulin
SEPT2
merge
GFP-Borg1
GFP-Borg2
GFP-Borg3

Fig. S2

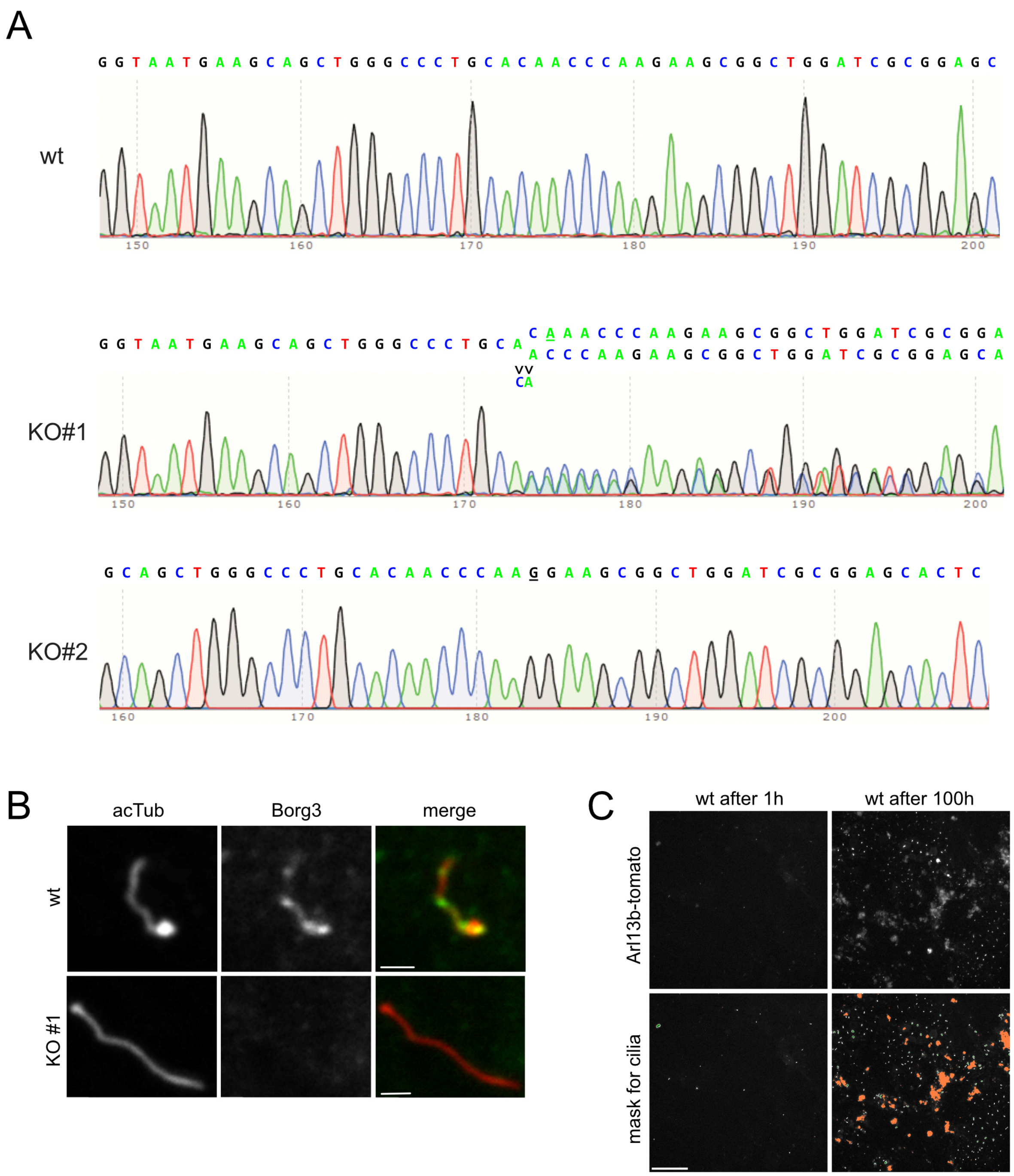
A
GGTAATGAAGCAGCTGGGCCCTGCACAACCCAAGAAGCGGCTGGATCGCGGAGC
wt
150
160
170
180
190
200
GGTAATGAAGCAGCTGGGCCCTGCA
CAAACCCAAGAAGCGGCTGGATCGCGGA
ACCCAAGAAGCGGCTGGATCGCGGAGCA
vv
CA
KO#1
150
160
170
180
190
200
GCAGCTGGGCCCTGCACAACCCAAGGAAGCGGCTGGATCGCGGAGCACTC
KO#2
160
170
180
190
200
B
acTub
Borg3
merge
wt
KO #1
C
wt after 1h
wt after 100h
Arl13b-tomato
mask for cilia

Fig. S3

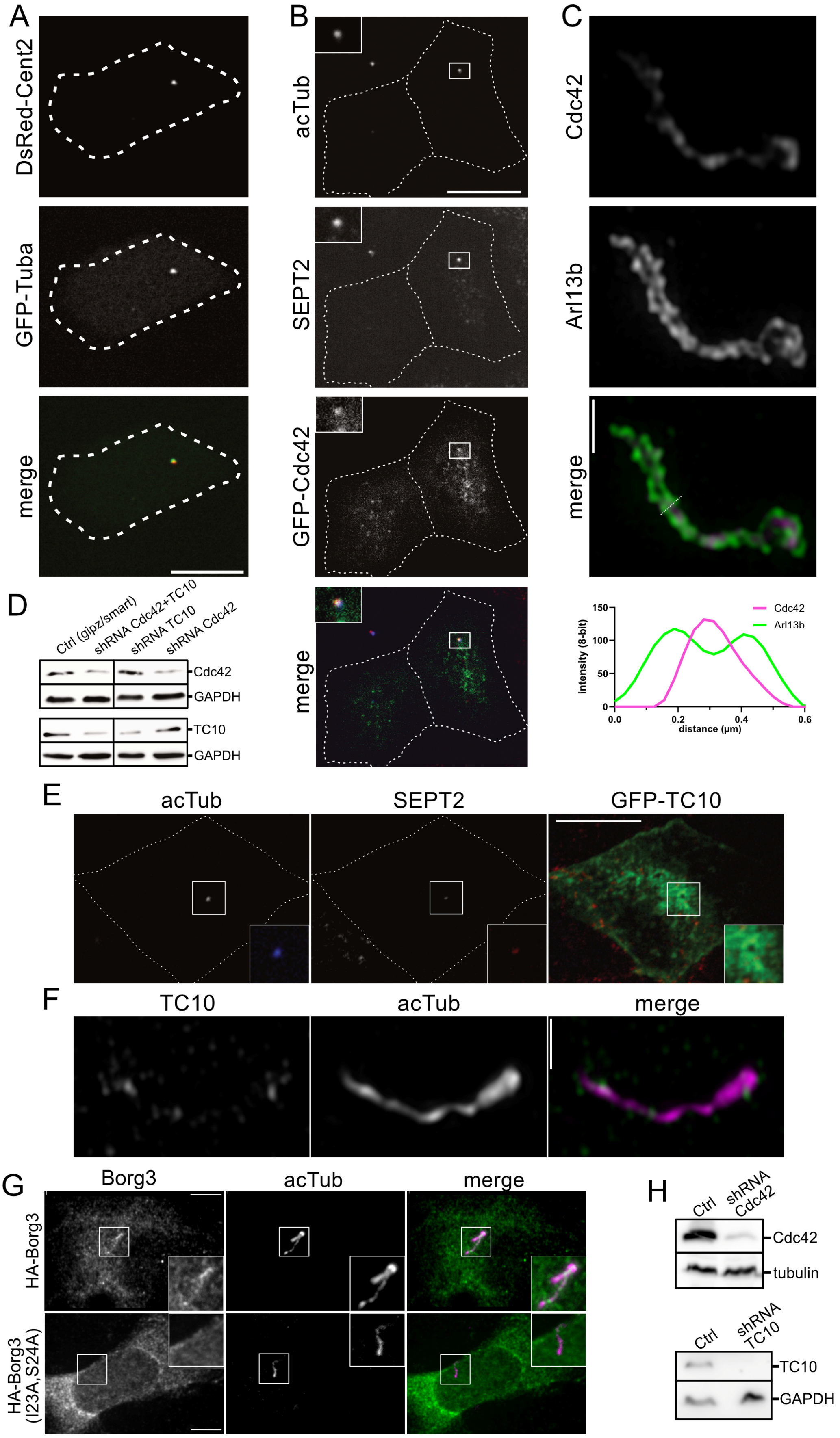
A
DsRed-Cent2
GFP-Tuba
merge
B
acTub
SEPT2
GFP-Cdc42
merge
C
Cdc42
Arl13b
merge
Cdc42
Arl13b
intensity (8-bit)
distance (µm)
D
Ctrl (gipz/smart)
shRNA Cdc42+TC10
shRNA TC10
shRNA Cdc42
Cdc42
GAPDH
TC10
GAPDH
E
acTub
SEPT2
GFP-TC10
F
TC10
acTub
merge
G
Borg3
acTub
merge
HA-Borg3
HA-Borg3 (I23A,S24A)
H
Ctrl
shRNA Cdc42
Cdc42
tubulin
Ctrl
shRNA TC10
TC10
GAPDH
